# Genomics of cellular proliferation under periodic stress

**DOI:** 10.1101/129163

**Authors:** Jérôme Salignon, Magali Richard, Etienne Fulcrand, Gaël Yvert

## Abstract

Living systems control cell growth dynamically by processing information from their environment. Although responses to one environmental change have been intensively studied, little is known about how cells react to fluctuating conditions. Here we address this question at the genomic scale by measuring the relative proliferation rate (fitness) of 3,568 yeast gene deletion mutants in out-of-equilibrium conditions: periodic oscillations between two salinity conditions. Fitness and its genetic variance largely depended on the stress period. Surprisingly, dozens of mutants displayed pronounced hyperproliferation at short periods, identifying unexpected controllers of growth under fast dynamics. We validated the implication of the high-affinity cAMP phosphodiesterase and of a regulator of protein translocation to mitochondria in this control. The results illustrate how natural selection acts on mutations in a fluctuating environment, highlighting unsuspected genetic vulnerabilities to periodic stress in molecular processes that are conserved across all eukaryotes.

## INTRODUCTION

Cells are dynamic systems that keep modifying themselves in response to variation of their environment. Interactions between internal dynamics of intracellular regulations and external dynamics of the environment can determine whether a cell dies, divides, differentiates or cooperates with other cells. For some systems, usually from model organisms, the molecules involved in signal transduction and cellular adaptation are largely known. Countless of them have been identified, often via genetic screens that isolated mutants with a defective response. How they act in motion, however, is unclear and it is difficult to predict which ones may be crucial upon certain frequencies of environmental fluctuations. In addition, since most screens were conducted in steady stressful conditions or after a single stress occurrence, molecules that are key to the dynamics may have been missed.

The control of cellular proliferation is essential to life and is therefore the focus of intense research, but its coupling to environmental dynamics remains poorly characterized. In addition, proliferation drives evolutionary selection, and the properties of natural selection in fluctuating environments are largely unknown. Although experimental data exist^1,2^, they are scarce and how mutations are selected in fluctuating conditions have mostly been studied under theoretical frameworks^3–6^. Repeated stimulations of a cellular response may have consequences on growth that largely differ from the consequences of a single stimulus. First, a small growth delay after the stimulus may be undetectable when applied only once, but can be highly significant when cumulated over multiple stimuli. Second, growth rate at a given time may depend on past environmental conditions that cells ‘remember’, and this memory can sometimes be transmitted to daughter cells^7^. These two features are well illustrated by the study of Razinkov *et al.*, who reported that protecting yeast GAL1 mRNA transcripts from their glucose-mediated degradation resulted in a growth delay that was negligible after one galactose-to-glucose change but significant over multiple changes^8^. This effect is due to short-term ‘memory’ of galactose exposures, which is mediated by GAL1 transcripts that are produced during the galactose condition and later compete for translation with transcripts of the CLN3 cyclin during the glucose condition. Other memorization effects were observed on bacteria during repeated lactose to glucose transitions, this time due to both short-term memory conferred by persistent gene expression and long-term memory conferred by protein stability^9^.

The yeast response to high concentrations of salt is one of the best studied mechanism of cellular adaptation. When extracellular salinity increases abruptly, cell-size immediately reduces and yeast triggers a large process of adaptation. The translation program^10,11^ and turnover of mRNAs^12^ are re-defined, calcium accumulates in the cytosol and activates the calcineurin pathway^13^, osmolarity sensors activate the High Osmolarity Glycerol MAPK pathway^13,14^, glycerol accumulates intracellularly as a harmless compensatory solute^14^, and membrane transporters extrude excessive ions^13^. Via this widespread adaptation, hundreds of genes are known to participate to growth control after a transition to high salt. What happens in the case of multiple osmolarity changes is less clear, but can be investigated by periodic stimulations of the adaptive response. For example, periodic transitions between 0 and 0.4M NaCl showed that MAPK activation was efficient and transient after each stress except in the range of ∼8 min periods, where sustained activation of the response severely hampered cell growth^15^. How genes involved in salt tolerance contribute to cell growth in specific dynamic regimes is unknown. If a protein participates to the late phase of adaptation its mutation may have a strong impact at large periods and no impact at short ones. It is also possible that mutations affecting growth in dynamic conditions have been missed by long-term adaptation screens. As mentioned above, a slight delay of the lag phase of adaptation may remain unnoticed after a single exposure, but its effect would likely cumulate over multiple exposures and be under strong selection in a periodic regime. Thus, even for a well-studied system such as yeast osmoadaptation, our molecular knowledge of cellular responses may be modest when dynamics are to be understood.

Although microfluidics enables powerful gene-centered investigations, its limited experimental throughput is not adapted to systematically search for genes involved in the dynamics of a cellular response. Identifying such genes can be done by applying stimulations to mutant cells periodically and testing if the effect of the mutation on proliferation is averaged over time. In other word, does fitness (proliferation rate relative to wild-type) of a mutant under periodic stress match the time-average of its fitness in each of the alternating condition? This problem of temporal heterogeneity is equivalent to the homogenization problem commonly encountered in physics for spatial heterogeneity, where microscopic heterogeneities in materials modify macroscopic properties such as their stiffness or conductivity^16^. If fitness is homogeneous (averaged over time), it implies that the effect of the mutation on the response occurs rapidly as compared to the frequency of environmental changes, that is does not affect the response lag phase and that the mutated gene is not involved in specific memory mechanisms. In contrast, fitness inhomogeneity (deviation from time-average expectation) is indicative of a role of the gene in the response dynamics.

In this study, we present a genomic screen that addresses this homogenization problem for thousands of gene deletion mutations in the context of the yeast salt response. The results reveal how selection of mutations can depend on environmental oscillations and identify molecular processes that unsuspectedly become major controllers of proliferation at short periods of repeated stress.

## RESULTS

### Genomic profiling of proliferation rates in steady and periodic salt stress

We measured experimentally the contribution of thousands of yeast genes on proliferation in two steady conditions of different salinity, and in an environment that periodically oscillated between the two conditions. We used a collection of yeast mutants where ∼5,000 non-essential genes have been individually deleted^17^. Since every mutant is barcoded by a synthetic DNA tag inserted in the genome, the relative abundance of each mutant in pooled cultures can be estimated by parallel sequencing of the barcodes (BAR-Seq)^18,19^. We set up an automated robotic platform to culture the pooled library by serial dilutions. Every 3 hours (average cell division time), populations of cells were transferred to a standard synthetic medium containing (S) or not (N) 0.2M NaCl. The culturing program was such that populations were either maintained in N, maintained in S, or exposed to alternating N and S conditions at periods of 6, 12, 18, 24 or 42 hours (Fig. 1A). Every regime was run in quadruplicates to account for biological and technical variability. Duration of the experiment was 3 days and populations were sampled every day. After data normalization and filtering we examined how relative proliferation rates compared between the periodic and the two steady environments.

**Fig. 1.**
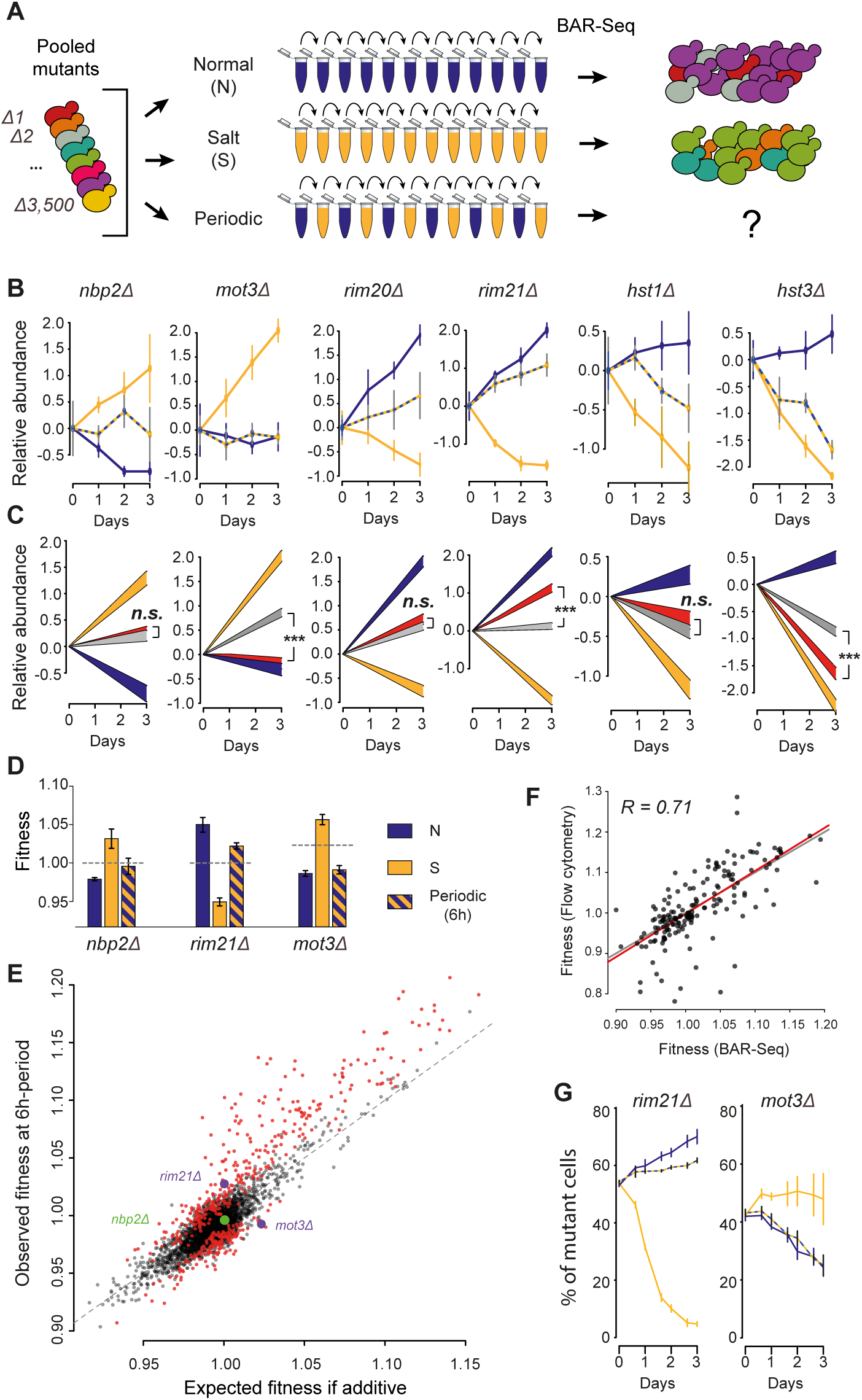
Genomic profiling of fitness in a periodic environment. (**A**) Experimental design. Populations of yeast deletion strains are cultured in media N (no salt), S (salt) and in conditions alternating between N and S at various periods. Allele frequencies are determined by BAR-seq and used to compute fitness (proliferation rate relative to wild-type) of each mutant. (**B**) Time-course of mutant abundance in the population, shown for six mutants. Relative abundance corresponds to the median of *log*_*2*_*(y/y*_*0*_) values ± *s.d.* (*n*=4 replicate cultures, except for condition N at day 3: *n*=3), where *y* is the normalized number of reads, and *y*_*0*_ is *y* at day 0. Conditions: N (yellow), S (blue), 6h-periodic (NS6, hatching). (**C**) Generalized linear models (*predicted value ± s.e.*) fitted to the data shown in (B), colored by condition: N (yellow); S (blue); NS6 predicted by the null model (grey) or predicted by the complete model including inhomogeneity (red). ***, *P* < 10^−8^. *n.s.*, non-significant. (**D**) Fitness values (w) computed from the data of two mutants shown in (B). Bars, mean ± *s.e.m.*, *n* = 3 (N) or 4 (S, NS6) replicate cultures, colored according to culture condition. Grey dashed line: expected fitness in case of additivity (geometric mean of fitness in N and S weighted by the time spent in each medium). (**E**) Scatterplot of all mutants showing their observed fitness under 6h-periodic fluctuations (*y*-axis, NS6 regime) and their expected fitness in case of additivity (*x*-axis, weighted geometric mean of fitness in N and S). Deviation from the diagonal reflects inhomogeneity. Red dots: 456 mutants with significant inhomogeneity (*FDR = 0.0001*, see methods). (**F**) Correlation between fitness estimates (w). Each dot corresponds to the median fitness of one mutant in one condition (N, S or NS6), measured from pooled cultures (*x*-axis) or from individual assays (one mutant co-cultured with WT cells, *y*-axis). Whole data: 52 mutants. *R*, Pearson coefficient; grey line, *y* = *x*; red line, linear regression. (**G**) Validation of inhomogeneity by cell counting. One graph shows the time-course of mutant abundance when it was individually co-cultured with GFP-tagged wild-type cells, measured by flow-cytometry. Median values ± *s.d.* (*n*=4 replicate cultures). Conditions: N (yellow), S (blue), 6h-periodic (NS6, hatching).

### Protective genes have diverse contributions to proliferation under periodic stress

We observed that genes involved in salt tolerance during steady conditions differed in the way they controlled growth under the periodic regime. As shown in Fig. 1B, differences were visible both among genes inhibiting growth and among genes promoting growth in high salt. For example, NBP2 is a negative regulator of the HOG pathway^20^ and MOT3 is a transcriptional regulator having diverse functions during osmotic stress^21,22^. Deletion of either of these genes improved tolerance to steady 0.2M NaCl (condition S). In the periodic regime, the relative growth of *mot3Δ* cells was similar to the steady condition N, as if transient exposures to the beneficial S condition had no positive effect. In contrast, the benefit of transient exposures was clearly visible for *nbp2Δ* cells. Differences were also apparent among protective genes. The Rim101 pathway has mostly been studied for its role during alkaline stress^13^, but it is also required for proper accumulation of the Ena1p transporter and efficient Na^+^ extrusion upon salt stress^23^. Eight genes of the pathway were covered by our experiment. Not surprisingly, gene deletion decreased (resp. increased) proliferation in S (resp. N) for all positive regulators of the pathway (Fig. 1B and Fig1-supplement-1). This is consistent with the need of a functional pathway in S and the cost of maintaining it in N where it is not required. The response to periodic stimulation was, however, different between mutants (Fig1-supplement-1). Although RIM21, DFG16 and RIM9 all code for units of the transmembrane sensing complex^24^, proliferation was high for *rim21Δ* and *dfg16Δ* cells but not for *rim9Δ* cells. Similarly, Rim8 and Rim20 both mediate the activation of the Rim101p transcriptional repressor^25,26^; but *rim8Δ* and *rim101Δ* deletions increased proliferation under periodic stress whereas *rim20Δ* did not. This pathway was not the only example displaying such differences. Cells lacking either the HST1 or the HST3 NAD(+)-dependent histone deacetylase^27^ grew poorly in S, but *hst1Δ* cells tolerated periodic stress better than *hst3Δ* cells (Fig. 1B).

Thus, gene deletion mutants of the same pathway or with similar fitness alterations in steady conditions can largely differ in their response to dynamic conditions.

### Widespread deviation from time-average fitness

We then systematically asked, for each of the 3,568 gene deletion mutants, whether its fitness in the periodic regime matched the time-average of its fitness in conditions N and S. We both tested the statistical significance and quantified the deviation from the time-average expectation. For statistical inference, we exploited the full BAR-seq count data, including all replicated populations, by fitting to the data a generalized linear model that included a non-additive term associated to the fluctuations (see methods). The models obtained for the six genes discussed above are shown in Fig. 1C. Overall, we estimated that deviation from time-average fitness was significant for as many as ∼2,000 genes, because it was significant for 2,497 genes at a False-Discovery Rate (FDR) of 0.2 (Supplementary Table 1). At a stringent FDR of 0.0001, we listed 456 gene deletions for which fitness inhomogeneity was highly significant.

For quantification, we computed fitness values as in Qian et al.^28^ (Fig. 1D) and plotted the observed fitness of all genes in the periodic environment as a function of their expected time-average fitness (Fig 1E). As for *nbp2Δ*, observed and expected values were often in good agreement. Highlighting the 456 significant genes revealed a surprising trend: for the majority of gene deletions expected to increase proliferation in the periodic regime (expected fitness > 1), observed fitness was unexpectedly high. Gene annotations corresponding to higher-than-expected fitness were enriched for transcriptional regulators and for members of the cAMP/PKA pathway (Supplementary Table 2), which is consistent with cellular responses to environmental dynamics.

Although BAR-Seq can estimate thousands of fitness values in parallel, it has two important limitations: estimation by sequencing is indirect and the individual fitness of a mutant is not distinguished from possible interactions with other mutants of the pool. We therefore sough to validate a subset of our observations by applying individual competition assays. Each mutant was co-cultured with a GFP-tagged wild-type strain, in N or S conditions or under the 6h-periodic regime, and the relative number of cells was counted by flow-cytometry^28,29^. Correlation between fitness estimates from BAR-Seq and individual assays was similar to previous reports^28,30^ (Fig. 1F, Fig1-supplement-2), and the assays unambiguously validated the fitness inhomogeneity of several mutants including *rim21Δ* and *mot3Δ* (Fig. 1G).

### Impact of environmental dynamics on mutants proliferation

If fitness inhomogeneity (deviation from time-average) is due to environmental dynamics, then it should be less pronounced at large periods of fluctuations. To see if this was the case, we computed for each mutant the ratio between its observed fitness in periodic stress and the time-average expectation from its fitness in the two steady conditions N and S. Fitness is inhomogeneous when this ratio deviates from 1. Plotting the distribution of this ratio at each period of fluctuation showed that, as expected, inhomogeneity was less and less pronounced as the period increased (Fig. 2A). We examined more closely three mutants displaying the highest inhomogeneity at the 6h period. Plotting their relative abundance in the different populations over the time of the experiment clearly showed that fitness of these mutants was unexpectedly extreme at short periods but less so at larger periods (Fig. 2B).

**Fig. 2.**
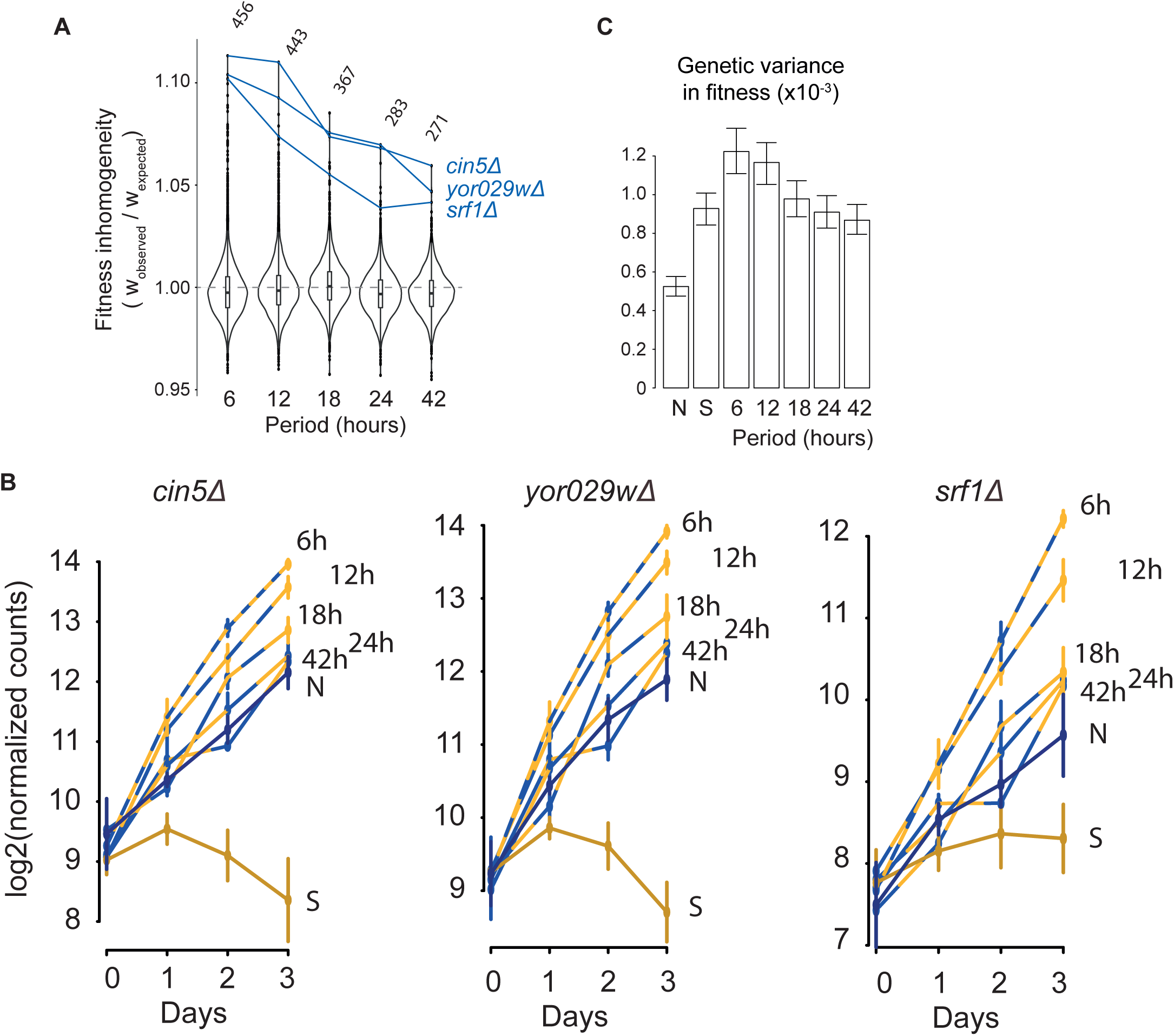
Proliferative advantage depends on environmental dynamics. (**A**) Violin plots showing the distribution of fitness inhomogeneity of 3,568 gene deletions at the indicated periods of environmental fluctuations. Traces and labels, mutants with extreme inhomogeneity at 6h-period. Top, number of gene deletions with significant inhomogeneity at *FDR = 0.0001*. (**B**) Time-course of the abundance of mutants *cin5Δ*, *srf1Δ* and *yor029w* in the pool of all mutants, under different fluctuating regimes, quantified by BAR-Seq. Median values ± *s.d.* (*n*=4 replicate cultures, except for the N condition at day 3: *n*=3). (**C**) The genetic variance in fitness of the pooled population of mutants was computed for each condition. Bars: 95% CI bootstrap intervals.

The fact that some mutants but not all were extremely fit to short-period fluctuations raised the possibility that the extent of differences in fitness between mutants may change with the period of environmental fluctuation. To see if this was the case, we computed the genetic variance in fitness of each pooled population of mutants (see methods). Fitness variation between strains was more pronounced when populations were grown in S than in N, which agrees with the known effect of stress on fitness differences^31^. Remarkably, differences were even larger in fast-fluctuating periodic regimes, but not slow-fluctuating ones (Fig. 2A). This shows that environmental fluctuations can exert additional selective pressures at the level of the whole population (see discussion).

### Fitness during alternating selection

Some gene deletions improved growth in one steady condition and penalized it in the other. This phenomenon is a special case of gene x environment interaction and is called antagonistic pleiotropy (AP)^28^. It is difficult to anticipate whether such mutations have a positive or negative impact on long-term growth in a periodic regime that alternates between favorable and unafavorable conditions, especially since fitness is not necessarily homogenized over time. We therefore studied these cases in more detail.

First, we examined if fitness inhomogeneity was related to the difference in fitness between the steady conditions (Fig. 3A). Interestingly, gene deletions conferring higher fitness in N than in S tended to be over-selected in the 6h-periodic regime, revealing a set of yeast genes that are costly in standard laboratory conditions as well as in the fast-fluctuating regime. We then searched for gene deletions that were advantageous in one steady condition and deleterious in the other (AP deletions). We found 48 gene deletions with statistically-significant AP between the N and S conditions (FDR = 0.01, Supplementary Table 3, see methods and Fig3-supplement-1). Interestingly, three of these genes coded for subunits of the chromatin-modifying Set1/COMPASS complex (Supplementary Table 2 and Fig3-supplement-2). We inspected whether the direction of effect of these 48 deletions depended on the period of fluctuations (Fig. 3B). For 33 (resp. 6) AP deletions, the effect was positive (resp. negative) at all periods. For two mutations (*vhr1Δ* and *rim21Δ*), the direction of selection changed with the oscillating period. To visualize the periodicity-dependence of all AP deletions, we clustered them according to their fitness inhomogeneity (Fig. 3C-D). This highlighted 5 different behaviours: fluctuations could strongly favour proliferation of a mutant at all periods (e.g. *cin5Δ*) or mainly when they were fast (e.g. *oca1Δ*), they could mildly increase (e.g. *rim101Δ*) or decrease it (e.g. *csf1Δ*) or they could both increase and decrease it depending on their period (*vhr1Δ*). Thus, fitness during alternating selection was generally asymmetric in favour of positive selection, and its dependency to the alternating period differed between genes.

**Fig. 3.**
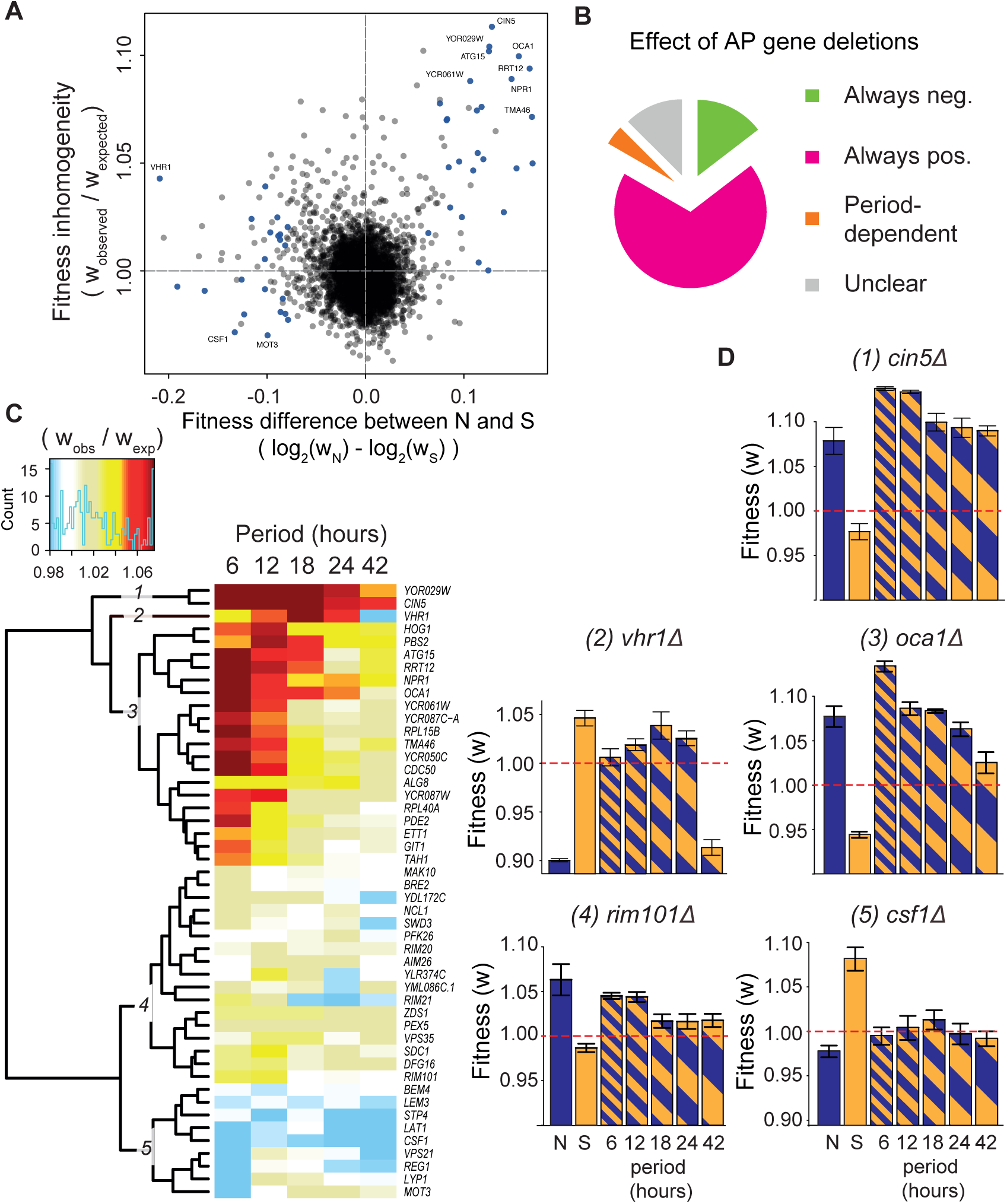
Long-term effect on growth during alternating selection. (**A**) Fitness inhomogeneity *vs.* antagonism between environments. Blue dots, 48 gene deletions with significant antagonistic pleiotropy (AP) between N and S (*FDR = 0.01*). (**B**) AP gene deletions were classified according to their direction of effect on growth, positive meaning advantageous. ‘Always’ means ‘at all periods of fluctuations’. (**C**) Hierarchical clustering of AP deletions according to fitness inhomogeneity. (**D**) Fitness values of five mutants representative of the clusters shown in C. Bars: mean ± *s.e.m.*, *n* = 3 (N) or 4 (others) replicate cultures.

### Environmental oscillations exacerbate the proliferation of some mutant cells

We made the surprising observation that fitness during fluctuations could exceed or fall below the fitness observed in both steady conditions (Fig. 2B), a behaviour called *’transgressivity’* hereafter. By using the available replicate fitness values, we detected 55 (resp. 23) gene deletions where fitness in the periodic environment was significantly stronger (resp. weaker) than the maximum (resp. minimum) of fitness in N and in S (Fig 4A, FDR=0.03, see methods). Importantly, transgressivity was observed not only from BAR-Seq but also when studying gene deletions one by one in competition assays, as shown for *pde2Δ, tom7Δ, trm1Δ* and *yjl135wΔ* (Fig. 4B-E). This reveals that environmental oscillations on short time scales can twist natural selection in favour of a subset of mutations on the long term. This may have important implications on the spectrum of mutations found in hyperproliferative clones that experienced repetitive stress (see discussion). It is also remarkable that the gene deletions displaying this effect were associated to various cellular and molecular processes: cAMP/PKA (*pde2Δ*), protein import into mitochondria (*tom7Δ*), autophagy (*atg15Δ*), tRNA modification (*trm1Δ*), phosphatidylcholine hydrolysis (*srf1Δ*) and MAPK signalling (*ssk1Δ*, *ssk2Δ*); and some of these molecular functions were not previously associated to salt stress.

**Fig. 4.**
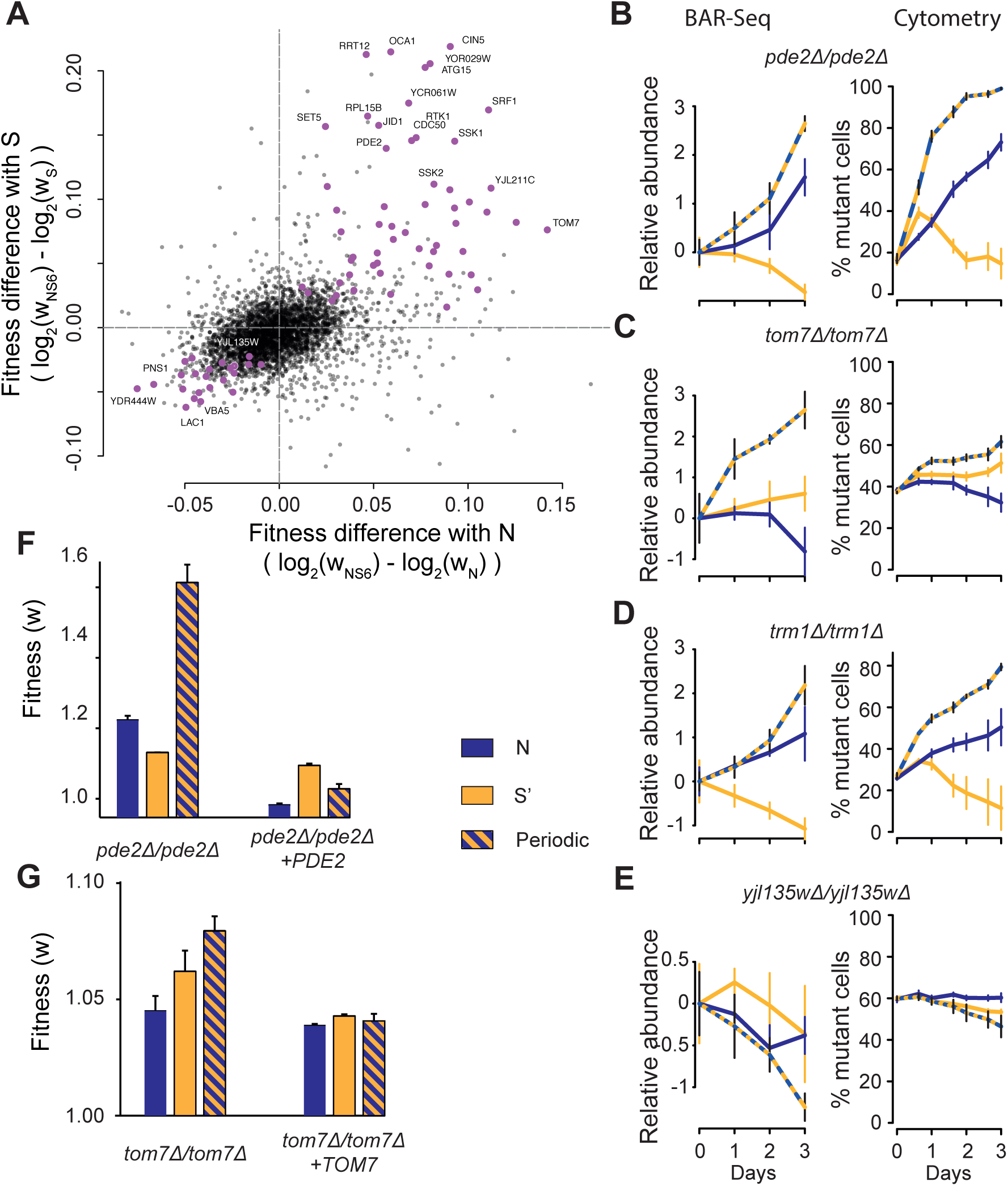
Extreme proliferation rates emerging from environmental oscillations. (**A**) Scatterplot of all mutants showing their observed fitness in the 6h-periodic regime (NS6) relative to their fitness in N (*x*-axis) and S (*y*-axis). Violet, 78 mutants with significant transgressivity (*FDR = 0.03*). (**B-E**) Time-course of mutant abundance in the pool of all mutants (BAR-Seq, left, as in Fig. 1B) or when the mutant was individually co-cultured with GFP-tagged wild-type cells (Flow-cytometry, right, as in Fig. 1G). Median values ± *s.d.* (*n*=4 replicate cultures, except for BAR-Seq N condition at day 3: *n*=3). Conditions: N (yellow), S (blue), NS6 (hatching). (**F-G**) Complementation assays. Diploid homozygous deletion mutants for *pde2* and *tom7* (strains GY1821 and GY1804, respectively) were complemented by integration of the wild-type gene at the *HO* locus (strains GY1929 and GY1921, respectively). Strains were co-cultured for 24h with GFP-tagged wild-type cells (strain GY1961) and relative fitness was measured by flow cytometry. Conditions: N (blue), S’ (0.4M NaCl; orange) and 6h-periodic fluctuations between N and S’ (hatching). Bars, mean fitness ± *s.e.m.* (*n*=3 replicate cultures).

### The high-affinity cAMP phosphodiesterase and Tom7p are necessary to limit hyperproliferation during periodic salt stress

As mentioned above, several gene deletions impairing the cAMP/PKA pathway displayed inhomogeneous fitness (Supplementary Table 2). One of them, *pde2Δ*, had a particularly marked fitness transgressivity (Fig. 4B). To determine if this effect truly resulted from the loss of PDE2 activity, and not from secondary mutations or perturbed regulations of neighboring genes at the locus, we performed a complementation assay. Re-inserting a wild-type copy of the gene at another genomic locus reduced hyperproliferation and fully abolished fitness transgressivity (Fig. 4F). Thus, the observed effect of *pde2Δ* directly results from the loss of Pde2p, the high-affinity phosphodiesterase that converts cAMP to AMP^32^, showing that proper cAMP levels are needed to limit proliferation during repeated salinity changes.

Unexpectedly, we found that deletion of TOM7, which has so far not been associated to saline stress, also caused fitness transgressivity in the 6h-periodic environment (Fig. 4C). The Tom7p protein regulates the biogenesis dynamics of the Translocase of Outer Membrane (TOM) complex, the major entry gate of cytosolic proteins into mitochondria^33^, by affecting both the maturation of the central protein Tom40p and the later addition of Tom22p^34,35^. We observed that re-inserting a single copy of TOM7 in the homozygous diploid mutant was enough to reduce hyperproliferation, although not to the levels of the wild-type diploid, and abolished fitness transgressivity (Fig. 4G). This suggests that proper dynamics of TOM assembly at the outer mitochondrial membrane are needed to limit proliferation during salinity fluctuations.

## DISCUSSION

We quantified the contribution of 3,568 yeast genes to cell growth during periodic salt stress. This survey showed that for about 2,000 genes, fitness was not homogenized over time. In other words, the observed fitness of these genes in periodic stress did not match the time-average of the fitness in the two alternating conditions. This widespread and sometimes extreme time-inhomogeneity of the genetic control of cell proliferation has several important implications.

### Novel information is obtained when studying adaptation out of equilibrium

A large part of information about the properties of a responsive system is hidden at steady state. For example, a high protein level does not distinguish between fast production and slow degradation. For this reason, engineers working on control theory commonly study complex systems by applying periodic stimulations, a way to explore the system’s behaviour out of equilibrium. Determining the frequencies at which a response is filtered or amplified is invaluable to predict the response to various types of stimulations. Such spectral analysis can sometimes reveal vulnerabilities, and it has also been applied to biological systems^36^. In the case of the yeast response to salt, Mitchell *et al.*^15^ monitored activation of the HOG pathway upon periodic stimulations and reported a resonance phenomenon at a bandwidth that was consistent with the known kinetics of the pathway.

In the present study, mutant cells used in a genomic screen were repeatedly stimulated by a periodic stress. This revealed two features of the salt stress response that were not suspected. Numerous gene deletions exacerbate hyperproliferation at short fluctuation periods (Figs. 2A and 4A); and many of the genes concerned were not previously associated to salt stress (e.g. TOM7, ATG15, SRF1. RPL15B, RRT12…). Thus, combining spectral analysis with genetic screening can reveal novel information on a well-known biological system.

### Gene x Environment interactions in dynamic conditions

Interactions between genes and environmental factors (GxE) are omnipresent in genetics and constitute the driving force for the adaptation of populations. Because model organisms offer the possibility to study a given genotype in various environmental conditions, they have been very useful to delimit the properties and extent of GxE. However, this has usually been done by comparing steady environments. Our observation that the dynamics of the environment can twist the effect of a mutation beyond what is observed in steady conditions raises a fundamental question: is GxE predictable when environmental dynamics are known? Since we observed unpredictable inhomogeneities mostly at short periods of environmental oscillations, the answer to this question likely depends on the speed of environmental fluctuations. It will therefore be helpful to determine what is the critical period below which prediction is challenging. We showed that for a given system (yeast tolerance to salt) this limit differed between mutations. Future experiments that track the growth of specific mutants in microfluidic chambers may reveal the bandwith of frequencies at which GxE interactions take place.

It is important to distinguish a periodic stress that is natural to an organism from a periodic stress that has never been experienced by the population (as considered here). In the first case, populations can evolve molecular clocks adapted to the stress period. This capacity is well known: nature is full of examples, and artificial clocks can be obtained by experimental evolution of micro-organisms^37^. In the case of periodic stress, an impressive result was obtained on nematodes evolving under anoxia/normoxia transitions at each generation time. An adaptive mechanism emerged whereby hermaphrodites produced more glycogen during normoxia, at the expense of glycerol that they themselves needed, and transmitted this costly glycogen to their eggs in anticipation to their need of it in the upcoming anoxia condition^38^. In contrast, when a periodic stress is encountered for the first time, cells face a novel challenge. The dynamic properties of their stress response can then generate extreme phenotypes, such as hyperproliferation, as described here (Fig. 2B, Fig. 4A-D), or long-term growth arrest as described by others^15^.

### Natural selection in fluctuating environments

Because the traits we quantified were the relative rates of proliferation between different genotypes (fitness), our survey provides a genome-scale view of natural selection during periodic stress. The impact of environmental fluctuations is a fundamental and complex subject, since natural environments and population adaptation are both dynamic. Population parameters such as allele frequencies, mutation rate, population size, target size for beneficial mutations determine the dynamics of genetic adaptation and they themselves depend on environmental conditions and therefore on environmental dynamics. Theoretical studies have shown that this complex interaction between the dynamics of adaptation and those of the environment can affect selection^3–5^. One of these studies modeled the fate of a *de novo* mutation appearing in a fixed-size population under a regime that fluctuated between two conditions, and causing a symmetric antagonistic effect between the two conditions^5^. The fluctuations were predicted to reduce the efficiency of selection in a way that, in addition to the fluctuating period, depended on two key factors i) the critical time necessary for a *de novo* mutation appearance and ii) the contrast in selection between the two conditions (equation [6] of Cvijovic *et al.*^5^). As a result, the effect of the mutation significantly deviated from the time-average of its effect in each of the conditions. It is important to distinguish this inhomogeneity from the one we describe here. First, we did not measure the effect of *de novo* mutations but of mutations that were all present prior to the fluctuations. Although rare additional mutations could arise afterwards, their effect would only be significant in the case of dominance (because we used homozygous diploid strains), and convergence (we monitored several replicate populations in parallel), which is very unlikely for less than 30 generations. Second, the mutations we studied did not necessarily have a symmetric effect between the two conditions (see *srf1Δ* in Fig. 2B for example). Conclusions of the two studies are therefore complementary: Cvijovic *et al.* reported a reduced selection on *de novo* mutations appearing during slow environmental fluctuations with seasonal drift, and we report here the emergence of strong positive selection on pre-existing mutations when novel, fast and strictly-periodic environmental fluctuations occur. These two types of inhomogeneity may both participate to the complexity of selection in natural environments.

Consistent with the inhomogeneities of fitness observed at the level of individual mutants, we also observed that the diversity of fitness among the pooled population of mutants was modified by environmental dynamics: the shorter the period of fluctuations, the stronger were the differences. This finding is important because, according to Fisher’s theorem, genetic variation in fitness reflects the rate of population adaptation^39,40^. Our observations therefore directly couple two time scales: fast dynamics at the level of environmental fluctuations with long-term changes of the population. Note that this link has been studied experimentally since the 1960’s: by evolving natural populations of *Drosophila* flies in either steady or fluctuating conditions, several studies showed that the genetic variance of fitness-related traits increased in the populations that evolved in fluctuating regimes^41–43^. In our case, the genetic diversity (a large pool of *de novo* mutations) pre-existed the fluctuations and the observed elevated genetic variance in fitness corresponds to a large diversity of selection coefficients (fitness itself) acting on the mutations when the environment fluctuates. Thus, the dynamics of natural environments may increase not only the genetic variance of fitness-related traits but also the diversity of the selection coefficients acting on mutations. Both of these effects would then participate to the coupling between the short time scales of environmental fluctuations and the long time scales of population adaptation.

### To sense, to memorize, or to anticipate ?

A mutation may improve fitness under periodic stress in several ways. It may render individuals highly sensitive and reactive to environmental changes, so that the lag following each change is reduced. A mutation may also modify the ability of cells to ‘remember’ past conditions. Yeast cells are known to ‘record’ stress occurrence via molecular changes conferring long-term (epigenetic) memory associated with an improved response at later exposures^44^. In the case of salt stress, this process involves chromatin modifications mediated by the Set1/COMPASS complex^45^. Mutants of this complex displayed a systematic fitness pattern in our data. Removal of either one of five components (Swd1p, Spp1p, Sdc1p, Swd3p, Bre2p) decreased fitness in N, increased it in S, and increased it similarly in the periodic regime (Fig3-supplement-2). This could result from memory alterations that change the response dynamics in ways that are better suited to the periodic regime. Alternatively, it could result from a trade-off: the benefits of epigenetic memory also have a cost. The mechanism consumes energy (remodelling), chemicals (e.g. AdoMet), and modifies chromatin instead of letting it free to replicate. This may penalize growth of wild-type cells if they do this repeatedly, as compared to mutants that do not. Stress memorization may also explain the fitness inhomogeneity of mutants impairing other chromatin modifying complexes, such as *rtt106Δ*, *set5Δ*, *swr1Δ, vps72Δ, hst3Δ* or *cac2Δ* (Supplementary Dataset). Finally, mutations may also diversify phenotypes between individual cells, or reduce the specialization of their phenotype, in anticipation of upcoming changes (bet-hedging)^46,47^. The relative efficiencies of these strategies and how they can evolve is a debated question^3,6^. Our screen offers new possibilities to investigate these adaptive strategies, for example by tracking the dynamics of growth of individual mutant cells in a controlled dynamic environment^15,48^. This may highlight genes that, when mutated, favour one strategy or the other.

### The high-affinity cAMP phosphodiesterase constitutes a genetic vulnerability to environmental dynamics

One of the mutants unexpectedly fit to stress oscillations was *pde2Δ*, and this phenotype was complemented by ectopic re-insertion of a wild-type copy of the gene. The yeast genome encodes two phosphodiesterases, one of low affinity that shares homology with only a fraction of eukaryotes (Pde1p), and one of high affinity that belongs to a well-studied class of phosphodiesterases found in many species, including mammals (Pde2p)^49^. We note that our genomic data did not indicate any obvious fitness alteration of *pde1Δ* cells in fluctuating conditions (Supplementary Dataset). These two enzymes convert cAMP into AMP. By binding to the Bcy1p repressor of Protein Kinase A, cAMP activates this complex and thereby promotes proliferation in optimal growth conditions. This regulation is implicated in the response to various stresses, including high salt^50,51^. Negative regulators of the pathway, including *PDE2*, are recurrent targets for *de novo* mutations in yeast populations evolving in steady experimental conditions^30^ and for natural standing variation affecting proliferation under stressful conditions^52^. The fitness transgressivity of *pde2Δ* cells that we observed suggests that the positive selection of such mutations may be even stronger if environmental conditions fluctuate. In addition, the output of the cAMP/PKA pathway is most likely governed by its dynamic properties, since intracellular levels of cAMP oscillate, with consequences on the stress response nucleo-cytoplasmic oscillations of Msn2p^53^. The activity of Pde2p is itself modulated by PKA^54^, and this negative feedback is probably important for suitable dynamics^55^. Our results suggest that loss of this feedback confers a hyperproliferative advantage and that it therefore constitutes a genetic vulnerability during prolonged exposure to periodic stress.

### Relevance to cancer

Cancer is an evolutionary issue: hyperproliferative cells possessing tumorigenic somatic mutations accumulate in tissues and threaten the life of the body. This process is driven by two main factors: occurrence of these mutations (mutational input) which depends both on the mutation rate and on the genomic target size of tumorigenesis, and natural selection of somatic mutations among cells of the body. The effect of mutations on proliferation rates is not the sole process of selection (tumors also evolve more complex phenotypes such as invasiveness or angiogenesis) but it is central to it; and human tissues are paced by various dynamics. Sleep, food intake, hormonal cycles, exercise, breathing, heart beats, circadian clocks, walking steps and seasons constitute a long list of natural rhythms, mechanic and electromagnetic waves as well as periodic medicine intake constitute artificial ones. To our knowledge, the impact of these dynamics on the selection process of somatic mutations has not been studied. Our results on yeast suggest that it may be significant, because a transient episode of periodic stress may strongly reshape allele frequencies in a population of mutant cells. If this happens in human tissues, it may affect the selection process of tumorigenic mutations. Also, if understood, such an effect could open medical perspectives to counter-select undesired mutations by applying beneficial environmental dynamics.

Remarkably, some of the yeast mutants displaying increased hyperproliferation during fast periodic stress correspond to molecular processes that are common to all eucaryotes (cAMP/PKA, autophagy, tRNA modifications, protein import in mitochondria). Our results suggest that the integrity of these pathways is threaten by environmental dynamics when wild-type yeast grow under periodic stress: if null mutations arise in the genes we identified, their high positive selection may cause their fixation. This raises the possibility that similar threats exist in humans: environmental dynamics may favour the loss of molecular functions that are important to limit proliferation. In particular, the RAS/cAMP/PKA pathway is altered in many cancers. Human cAMP-phosphodiesterases have been associated to tumor progression both positively (PDEs being overexpressed in tumors and PDE inhibitors limiting proliferation in several contexts^56^) and negatively (predisposing mutations being found in PDE8B^57^ and PDE11A^58–60^). Given our observations, it is possible that the dynamics of the cellular environment may modulate the effect of these deregulations. More generally, now that barcoding techniques allow to track selection in cancer cell lines^61^, using them in a context of periodic stimulations may reveal unsuspected genetic factors.

## METHODS

### Yeast deletion library and growth media

The pooled homozygous diploid Yeast Deletion Library was purchased from Invitrogen (ref. 95401.H1Pool). In each strain, the coding sequence of one gene had been replaced by a KanMX4 cassette and two unique barcodes (uptag and downtag) flanked by universal primers ^62^. Following delivery, the yeast pool was grown overnight in 100ml YPD medium, and 500 μl aliquots (2.2x10^8^ cells/ml) were stored in 25% glycerol at -80°C. Medium N (Normal) was a synthetic complete medium made of 20 g/L D-glucose, 6.7 g/L Yeast Nitrogen Base without amino-acids (Difco), 88.9 mg/L uracil, 44.4 mg/L adenine, 177.8 mg/L leucine and all other amino-acids at 88.9 mg/L and 170 μl/L NaOH 10N. Medium S (Salt) was made by adding 40 ml/L NaCl 5M to medium N (final concentration of 0.2M).

### Fluctuation experiment setup

All steps of the fluctuation experiment were carried out in 96-well sterile microplates using a Freedom EVO200 liquid handler (Tecan) equipped with a 96-channel pipetting head (MCA), a high precision 8-channel pipetting arm (LiHa), a robotic manipulator arm (RoMa), a Sunrise plate reader (Tecan), a MOI-6 incubator (Tecan), and a vacuum station (Millipore). All robotic steps were programmed in Evoware v2.5.4.0 (Tecan). Each of 7 culture conditions (N, S, NS6, NS12, NS18, NS24, NS42) was applied on four independent populations. To reduce technical variability and population bottlenecks, each population was dispatched in four parallel microplates before each incubation step and these plates were combined into a single one after incubation. The size of each population was maintained over 2.1 x 10^7^ cells.

### Initialization of pooled-mutants cultures

Four aliquots of the yeast deletion library were thawed, pooled and immediately diluted into 100 ml of fresh N medium. After mixing, samples of 220 μl of the cell suspension were immediately distributed into 28 wells of each of four distinct microplates. This initiated a total of 112 populations of cells, each containing ∼320 copies of each mutant strain on average. Plates were then incubated at 30°C for 6 hours with 270 rpm shaking.

### Fluctuations of pooled-mutants cultures

Twice a day, a stock of source plates that contained sterile N or S fresh medium in the appropriate wells were prepared. Every 3 hours, the four microplates containing cells were removed from the incubator (30°C, 270 rpm) and cells were transferred to a single sterile plate having a 1.2 μm-pore filter bottom (Millipore, MSBVS1210), media were removed by aspiration, four fresh source plates were extracted from the stock, 62 μl of sterile media was pipeted from each of them and transferred to the filter plate, cells were resuspended by pipetting 220 μl up and down, and 60 μl of cell suspension were transferred to each of the 4 source plates which were then incubated at 30°C with 270 rpm for another 3 hours. Every 6 hours, cell density was monitored for one of the four replicate plates by OD_600_ absorbance. Every 24 hours, 120 μl of cultures from each replicate plate were sampled, pooled in a single microplate, centrifuged 10 minutes at 5000g and cell pellets were frozen at -80°C. Dilution rates of the populations were: 85% when the action was only to replace the media, 55% when it was to replace the media and to measure OD, and 32% when it was to replace the media, to measure OD and to store samples.

The experiment lasted 78 hours in total and generated samples from 28 independent populations at time points 6h (end of initialization), 30h, 54h and 78h.

### BAR-seq

Frozen yeast pellets were resuspended in 200 μl of a mix of 30 ml of Y1 Buffer (91.1 g of sorbitol in 300 ml H_2_O, 100 ml of 0.5 M EDTA, 0.5 ml of β-mercaptoethanol, completed with 500 mL of water), 60 units of zymolyase (MPBiomedicals, ref 8320921) and 22.5 μl of RNAseA at 34 mg/ml (Sigma ref R4642), vortexed and incubated for 1 hour at 37°C for cell wall digestion. Genomic DNA (gDNA) was extracted by using the Macherey Nagel 96-well Nucleospin kit (ref 740741.24) following manufacturer’s instructions. We designed and ordered from Eurogentec a set of 112 reverse primers of the form 5’-P5-X_9_-U2-3’, where P5 (5’-AATGATACGGCGACCACCGAGATCTACACTCTTTCCCTACACGACGCTCTTCCGA TCT-3’) allowed Illumina sequencing, X_9_ was a custom index of 9 nucleotides allowing multiplexing via a Hamming code ^63^, and U2 (5’-GTCGACCTGCAGCGTACG-3’) matched a universal tag located downstream the uptag barcode of each mutant yeast strain. PCR amplification of the barcodes of each sample was done by using these reverse primers in combination with one forward primer of the form 5’-P7-U1-3’, where P7 (5’-CAAGCAGAAGACGGCATACGAGATGTGACTGGAGTTCAGACGTGTGCTCTTCCG ATCT-3’) allowed Illumina sequencing and U1 (5’-GATGTCCACGAGGTCTCT-3’) matched a universal tag located upstream the uptag barcode of each yeast mutant. Reagents used for one PCR reaction were: 18.3 μl of water, 6 μl of Buffer HF5X and 0.2 μl of Phusion polymerase (ThermoFischer Scientific, ref F530-L), 2.5 μl of dNTP 2.5 mM, 1 μl of each primer at 333 nM and 1 μl of gDNA at 300 to 400 ng/μl. Annealing temperature was 52°C, extension time 30 sec, and 30 cycles were performed. As observed previously, the PCR product migrated as two bands on agarose gels, which can be explained by heteroduplexes^64^. Both bands were extracted from the gel, purified and eluted in 30 μl water. All 112 amplification products were pooled together (10 μl of each), gel-purified and eluted in a final volume of 30 μl water and sequenced by 50nt single reads on a Illumina HiSeq2500 sequencer by ViroScan3D/ProfileXpert (Lyon, France).

### Data extraction, filtering and normalization

Demultiplexing was done via an error-correction Hamming code as described previously ^63^. Mapping (assignment of reads to yeast mutant barcodes) was done by allowing a maximal Levenstein distance of 1 between a read and any sequence in the corrected list of mutant barcodes of Smith et al. ^18^. In total, 291 million reads were mapped and used to build a raw 6,004 (mutants) by 112 (samples) table of counts. One sample was discarded because it was covered by less than 300,000 total counts and displayed mutants frequencies that were poorly correlated with their relevant replicates. Similarly, 2,436 mutants were covered by few (< 2,000) counts over all samples (including samples of another unrelated experiment that was sequenced in parallel) and were discarded, leaving a count table of 3,568 mutants by 111 samples for further analysis. This table was then normalized using the function *varianceStabilizingTransformation* from the DESeq2 package ^65^ (version 1.8.1) with arguments blind = FALSE and fitType = ‘local’.

### Fitness estimation

We followed the method of Qian *et al.* ^28^ to estimate the fitness cost or gain (*w*) of each mutant in each population. Eleven genes (Supplementary Table 4) were considered to be pseudogenes or genes with no effect on growth, and the data from the corresponding deletion mutants were combined and used as an artificial “wild-type” reference. For each mutant strain *M*, *w* was calculated as:

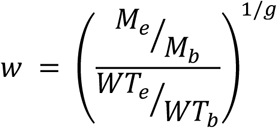

with *M*_*b*_, *M*_*e*_, *WT*_*b*_, and *WT*_*e*_ being the frequencies of strain *M* and artificial wild type strain (*WT*) at the beginning (*b*) or end (*e*) of the experiment, and *g* the number of generations in between. *g* was estimated from optical densities at 600nm of the entire population. It poorly differed between conditions and we fixed *g* = 24 (8 generations per day, doubling time of 3h).

### Deviation from time-average fitness

We analyzed fitness inhomogeneity by both quantifying it and testing against the null hypothesis of additivity. The quantification was done by computing 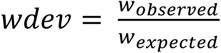, where *w*_*observed*_ was the fitness of the mutant strain experimentally measured in the periodic environment and *w*_*expected*_ was the fitness expected given the fitness of the mutant strain in the two steady environments (N and S), calculated as

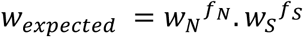

with *f*_*N*_ and *f*_*S*_ being the fraction of time spent in N and S media, respectively, during the course of the fluctuation experiment. Statistical inference was based on a Generalized Linear Model applied to the normalized count data. We assumed that the normalized counts of mutant *i* in condition *c* (N, S or periodic) at day *d* in replicate population *r* originated from a negative binomial distribution NB(*λ*_*i*_, *α*), with:

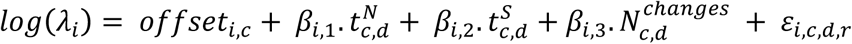

and *offset*_*i,c*_ being the median of normalized counts for condition *c* at day 0, 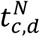, 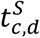 being the amount of time spent in medium N and medium S at day *d*, respectively, 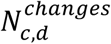 being the number of changes between the two media that took place between days 0 and *d*, and ε being the residual error. The model was implemented in R using the function *glm.nb* of the MASS package (version 7.3-40).

If fitness is homogenized in a fluctuating environment, then it is insensitive to the number of changes and *β*_*i,3*_ = 0. Inhomogeneity can therefore be inferred from the statistical significance of the term 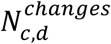 of the model. The corresponding *p*-values were converted to *q*-values, using package *qvalue* version 2.0.0 in order to control the False Discovery Rate.

**Genetic variance in fitness** was computed for each condition as:

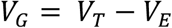

where

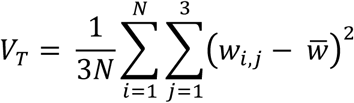

was the total variance, and

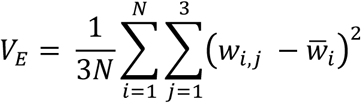

was an estimate of the non-genetic variance in fitness, with *N* being the number of gene deletions, *w*_*i,j*_ the fitness of gene deletion *i* in replicate *j*, 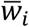 the mean fitness of gene deletion *i* and 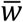 the global mean fitness. The 95% confidence intervals of V_G_ were computed from 1,000 bootstrap samples (randomly picking mutant strains, with replacement).

### Antagonistic Pleiotropy

We used the observed *w*_*N*_ and *w*_*S*_ values (fitness in the N and S steady conditions, respectively) of the deletion mutants to determine if a mutation was antagonistically pleiotropic (AP). Our experiment provided, for each mutant, 3 independent estimates of *w*_*N*_ and 4 independent estimates of *w*_*S*_ (replicate populations). For each mutant, we combined these estimates in 3 pairs of (*w*_*N*_, *w*_*S*_) values by randomly discarding one of the 4 available *w*_*S*_ values, and these pairs were considered as 3 independent observations. We considered that an observation supported AP if the fitness values (*w*_*N*_, *w*_*S*_) showed (1) an advantage in one of the conditions and a disadvantage in the other, and (2) deviation from the distribution of observed values in all mutants, since most deletions are not supposed to be AP. Condition (1) corresponded to: (*w*_*N*_ > 1 AND *w*_*S*_ < 1) OR (*w*_*N*_ < 1 AND *w*_*S*_ > 1). Condition (2) was tested by fitting a bivariate Gaussian to all observed (*w*_*N*_, *w*_*S*_) pairs and labelling those falling 2 standard deviations away from the model (Fig3-supplement-1). A deletion was considered AP if all 3 observations supported AP, which was the case for 48 deletions. A permutation test (re-assigning observations to different deletions replicates) determined that less than one deletion (0.54 on average) was expected to have three observations supporting AP by chance only (Supplementary Table 3). For the selected 48 deletions, the magnitude of AP was computed as *w*_*N*_ / *w*_*S*_. For each deletion, the direction of selection (Fig. 3C) in each condition was considered to be positive if 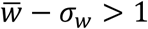, negative if 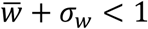 and ambiguous otherwise, with 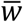 and *σ*_*w*_ being the mean and standard deviation of fitness values across replicates, respectively. A mutation was classified as: ‘unclear’ if its direction of selection was ambiguous at four or five fluctuation periods, ‘always positive’ (resp. ‘always negative’) if all its unambiguous directions of selection were positive (resp. negative) and ‘period-dependent’ if its direction differed between periods.

### Transgressive fitness

We considered that a mutant had transgressive fitness if at least 3 of its 4 observed replicate measures of fitness in fluctuating conditions (*w*_*NS*_) were either all higher than 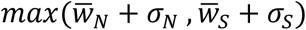 or all lower than 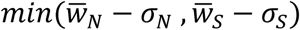, where 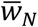 (resp. 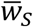) was the mean fitness value in steady condition N and S, respectively, and (resp. *σ*_*s*_) the corresponding standard deviation. A permutation test (re-assigning observations to random mutants) determined that less than three mutants (2.24 on average) were expected to display three replicates supporting transgressivity by chance only (Supplementary Table 5).

### Direct fitness measurement by flow cytometry: plasmids and strains

Individual homozygous diploid knock-out strains were ordered from Euroscarf. Oligonucleotides and modified strains used in this study are listed in Supplementary Tables 6 and 7, respectively. Wild-type strain BY4743 and individual mutants of interest were ordered from Euroscarf. We constructed a GFP-tagged wild-type strain (GY1738), and its non-GFP control (GY1735), by tranforming BY4743 with plasmids pGY248 and HO-poly-KanMX4-HO ^66^, respectively. Plasmid pGY248 was ordered from GeneCust who synthesized a Pact1-yEGFP BamHI fragment and cloned it into HO-poly-KanMX4-HO. Complemented strains were generated by cloning the wild-type copy of each gene of interest into a plasmid targeting integration at the *HO* locus. We first prepared a vector (pGY434) by removing the repeated *hisG* sequence of plasmid HO-hisG-URA3-hisG-poly-HO ^66^ by SmaI digestion and religation followed by ClaI digestion and religation. For *PDE2*, the wild-type (S288c) coding sequence with its 600bp upstream and 400bp downstream regions was synthesized by GeneCust and cloned in the BglII site of pGY434. The resulting plasmid (pGY453) was digested with NotI and transformed in strain GY1821 to give GY1929. For *TOM7*, we constructed plasmid pGY438 by amplifying the HOL-URA3-HOR fragment of pGY434 with primers 1O21 and 1O22, and cloning it into pRS315 ^67^ (linearized at NotI) by *in vivo* recombination. The wild-type copy of *TOM7* (coding sequence with its 465bp upstream and 813bp downstream regions) was PCR-amplified from strain BY4742 using primers 1O27 and 1O28 and co-transformed in BY4742 with PacI-PmeI fragment of pGY438 for *in vivo* recombination. The resulting plasmid (pGY442) was digested by NotI and the 4-kb fragment containing HO-URA3-TOM7-HO was gel-purified and transformed in GY1804 to obtain GY1921. Proper integration at the HO locus was verified by PCR. Since complementation was accompanied by the URA3 marker, which likely contributes to fitness, we competed strains GY1921 and GY1929 with a URA+ wild-type strain (GY1961), which was obtained by transforming strain GY1738 with the PCR-amplified URA3 gene of BY4716 (with primers 1D11 and 1D12). The non-GFP control URA+ wild-type strain GY1958 was obtained similarly.

### Direct fitness measurement by flow cytometry: fluctuation cultures

Each plate contained 8 different mixed cultures (one per row) and 3 different conditions (N, S, NS6) with 4 replicates each that were randomized (neighboring columns contained different conditions). Four plates were handled in parallel, which allowed us to test 32 different co-cultures per run, with at least one row per plate dedicated to controls (Wild-Type strain vs. itself or wild-type strain alone). Strains were streaked on G418-containing plates. Single colonies were used to inoculate 5 ml of N medium and were grown overnight at 30°C with 220 rpm shaking. The next day, concentration of each culture was adjusted to an OD_600_ of 0.2. For co-cultures, 2 ml of wild-type cell suspension was mixed with 2 ml of mutant cell suspension, and 220 μl of this mix was transfered to the desired wells of a microplate. Plates were then incubated on the robotic platform at 30°C with 270 rpm for 4-5h. Fluctuations of the medium condition were also done by robotics: dilution (keeping 130μl of the 220μl cell suspension), filtration and refill every 3 hours, using a stock of fresh source plates prepared in advance. Twice a day, 90 μl of the cell suspension were fixed and processed for flow-cytometry. Fixation was done on the robotic platform, by washing cells twice with PBS 1X, resuspending them in PBS 1X + Paraformaldehyde 2% and incubating at room temperature for 8 min, washing with PBS 1X, resuspending cells in PBS + Glycine 0.1M, incubating at room temperature for 12 min, and finally washing cells with PBS 1X and re-suspending them in PBS 1X. Plates were then diluted (at 80-95%) in PBS 1X and stored at 4°C before being analyzed on a FacsCalibur flow cytometer (BD Biosciences). Acquisitions were stored on 10,000 cells at a mean rate of 1,000 cells/s.

### Direct fitness measurement by flow cytometry: data analysis

Raw .fcs files were analyzed using the *flowCore* package (version 1.34.3) from Bioconductor ^68^ and custom codes. Cells of homogeneous size were dynamically gated as follows: (i) removal of samples containing less than 2000 cells, (ii) removal of events with saturated signals (FSC, SSC or FL1 ≥ 1023 or ≤ 0), (iii) computation of a density kernel of FSC,SSC values to define a perimeter of peak density containing 40% of events and (iv) cell gating using this perimeter, keeping >4,000 cells. In order to classify each cell as GFP^+^ or GFP^-^, FL1 thresholds were determined automatically using the function *findValleys* from package *quantmod* (version 0.4-4). The relevance of these thresholds was then verified on control samples containing only one of the two strains (unimodal GFP+ or GFP-). After classifying GFP^+^ (i.e. WT) and GFP^-^ (i.e. mutant) cells, fitness values were computed as 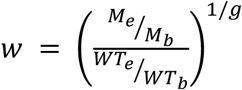, with *M*_*b*_, *M*_*e*_, *WT*_*b*_, and *WT*_*e*_ being the frequencies of mutant strain *M* and wild type strain (*WT*) at the beginning (*b*) or end (*e*) of the experiment, and *g=24* the number of generations in between.

## ACKNOWLEDGEMENTS

We thank Julien Gagneur for suggestions on normalization and general linear models, Arnaud Bonnaffoux, Florent Chuffart, Pascal Hersen, Abderrahman Khila, Sébastien Lemaire, Serge Pelet and Alexandre Soulard for discussions, Julien Gagneur, Jun-Yi Leu, Stephen Proulx, Mark Siegal and Henrique Teotonio for critical reading of the manuscript, Audrey Barthelaix for initial tests on the robotic platform, Hélène Duplus-Bottin for technical help on strains constructions, David Stillman for plasmids, Sandrine Mouradian and SFR Biosciences Gerland-Lyon Sud (UMS3444/US8) for access to flow cytometers and technical assistance, BioSyL Federation and Ecofect Labex for inspiring scientific events, developers of R, Bioconductor and Ubuntu for their software. This work was supported by the European Research Council under the European Union’s Seventh Framework Programme FP7/2007-2013 Grant Agreement n°281359 and by the Fondation ARC pour la recherche sur le cancer.

### AUTHOR CONTRIBUTIONS

J.S. and M.R. set up automated cultures; J.S. performed the experiments, optimized automation and analyzed the data; M.R. designed multiplexing oligonucleotides, set up BAR-Seq libraries preparations and supervised J.S. for the genomic experiment; J.S, M.R., and E.F. constructed strains; J.S. and E.F. performed flow cytometry; J.S., M.R., and G.Y. implemented the GLM model, interpreted results and wrote the paper; G.Y. conceived, designed and supervised the study.

## LIST OF SUPPLEMENTARY MATERIALS

**Figure1-figure-supplement 1. BAR-seq fitness profile of mutants of the Rim101 pathway.** (**A**) For mutants of the Rim101 pathway available in our data is shown their time-course abundance (left) and their fitted Generalized linear models (right), as in Figure 1. (**B**) Schematic representation of the pathway with colors corresponding to the level of fitness inhomogeneity of each member.

**Figure1-figure-supplement 2.** Time-course of mutant abundance, for mutants analyzed by BAR-Seq and individual competition assays.

**Figure3-figure-supplement 1. Detection of Antagonistic Pleiotropy.** Every dot corresponds to one mutant. Coordinates correspond to median fitness values of replicate populations grown in N (*n=3*) or S (*n=4*) condition. Oblique line: y=x. Red, AP mutants.

**Figure3-figure-supplement 2. BAR-seq fitness profile of mutants of the Set1/COMPASS complex.** (**A**) For each mutant of the complex available in our data is shown their time-course abundance (left) and their fitted Generalized Linear Model (right), as in Figure 1. (**B**) Schematic representation of the Compass complex (based on Soares *et al.*^69^) with colors corresponding to the level of fitness inhomogeneity of each member of the complex.

**Supplementary Table 1.** Number of deletion mutants having significant fitness inhomogeneity in the 6h-periodic regime, based on Generalized Linear Model.

**Supplementary Table 2.** Gene Ontology analysis.

**Supplementary Table 3.** Number of Antagonistic Pleiotropic mutants detected at various stringency.

**Supplementary Table 4.** Deletions of genes and pseudogenes used to infer Wild-Type fitness.

**Supplementary Table 5.** Number of mutants showing transgressive fitness at 6h-period fluctuations.

**Supplementary Table 6.** DNA primers used in this study.

**Supplementary Table 7.** Yeast strains used in this study.

**Supplementary Dataset.** Full genomic dataset. See README.txt file for documentation.

